# History bias and its perturbation of the stimulus representation in the macaque prefrontal cortex

**DOI:** 10.1101/2024.10.01.616011

**Authors:** Danilo Benozzo, Lorenzo Ferrucci, Francesco Ceccarelli, Aldo Genovesio

## Abstract

Multiple history biases affect our representation of magnitudes, such as time, distance, and size. It is not clear whether the previous stimuli interfere with the discrimination process from the moment of stimulus presentation, during working memory retention, or even later during the decision-making phase. We used a spatial discrimination task involving two stimuli of different magnitudes, presented sequentially at various distances from the center. The monkey’s task was to select the farthest of them. We showed that the previous stimulus magnitude generated a contraction bias effect, but only when its stimulus features differed from those of the current stimulus. In this case, at the neural level we also observed that the decoding of the stimulus magnitude achieved the highest accuracy when it matched the magnitude of the preceding stimulus for which the decoder was trained. This indicates that past stimuli can affect magnitude processing already during the stimulus presentation, even before the decision process. Interestingly, this effect manifested when the trace of the previous stimulus magnitude reactivated in the second part of the stimulus presentation after an “activity-silent” period.

## Introduction

Previous experience influences and biases our behavior. One of these biases is the central tendency bias, first reported by Hollingworth (Hollingworth et al., 1910), which is also known as the contraction bias. This bias involves the tendency for perceptual estimates to regress toward the mean of the distribution of the set of previously presented stimuli. Evidence of this bias has been described across various domains: in the auditory system (Raviv et al., 2014; Akrami et al., 2018), the visual system (Olkkonen et al., 2014), and the somatosensory system (Fassihi et al., 2017). Additionally, it has also been observed in the timing domain (Zhang and Zhou, 2017; Shi and Burr, 2016; Jazayeri and Shadlen, 2010; Roach et al., 2017; Cicchini et al., 2012).

In a controlled laboratory environment, where experimental trials are independent and often predictable, integrating past experiences with current information can introduce a contraction bias, leading to errors influenced by this bias. However, when viewed through a Bayesian framework, this contraction bias reveals its advantages, particularly in mitigating the effects of noisy data (Ashourian et al., 2011; Zhang and Zhou, 2017; Bausenhart et al., 2016; Olkkonen et al., 2014; Raviv et al., 2012; Petzschner et al., 2015).

Contraction bias combines its effect with other serial biases, all originating from previous information’s influence on the current trial. Their interconnection is still not completely understood, making it challenging to distinguish the individual contributions of each factor to the overall outcome. Recently, Boboeva et al., (2024) suggested a new perspective, proposing that contraction bias is an emergent effect of the interplay between history biases and working memory processing.

An explicit coding of previous information or of its impact on neural activity during the current trial is indicative of the effect of previous trials on performance. Several studies have shown the effect of trial history on current neural activity, particularly in relation to previous outcomes and behavioral goals (Barraclough et al., 2004; Genovesio et al., 2005; Seo et al., 2007; Seo and Lee, 2009_;_ Histed et al., 2009; Genovesio et al., 2014; Hermoso-Mendizabal et al., 2020). Other studies have shown that past experience can not only be represented at the neural level but also bias future choices in the prefrontal cortex, a phenomenon that has been less extensively studied (Padoa-Schioppa 2013; Barbosa et al., 2020; Mochol et al., 2021).

While previous studies have shown that information from previous trials keeps getting encoded in the next trial, it remains unclear whether previous trials can influence the coding of stimulus magnitude information during the stimulus presentation as well, possibly leading to behavioral biases.

Only a few studies investigated the neural correlates of contraction bias: Akrami et al. (2018) in the rat’s parietal cortex and Benozzo et al. (2023) in the monkey’s prefrontal cortex. Through optogenetic inactivation, Akrami et al. (2018) showed that the effect of the previous trial was eliminated by switching off the rat parietal cortex neurons. This translated into an improvement in behavioral performance, showing that, together with their neurophysiological results, at least in rats, the parietal cortex combines current and past information.

We used a two-stimuli distance discrimination task to investigate whether and how the most recent trial interferes with the processing of the current trial stimuli. In principle, information from the previous trial could bias perceptual decisions during any of three task phases: first, during stimulus presentation; second, during the maintenance of the stimulus in memory; or third, during the comparison process when the first and second stimuli are compared. We have previously shown that contraction bias affects the decision process (Benozzo et al., 2023) by biasing towards the average distribution of stimulus magnitude. However, this did not rule out the possibility that the influence of prior information begins even earlier in the task. In this study, we investigated whether the most recent experience, that is the last stimulus presented in the previous trial, can also affect the representation of the current stimulus, thus placing the effect of the previous trial bias to an earlier stage than the decision process.

We found that the decoding of the magnitude of the stimulus to be discriminated could be discriminated with higher accuracy when training and testing were performed using the same previous stimulus magnitudes, pointing to an effect of the past stimulus magnitude on the processing of the stimuli. This effect was not strictly time-locked, though it tended to emerge more strongly in the second part of the stimulus presentation, coinciding with the reactivation of the trace of the past stimulus magnitude, after it was not explicitly coded in the initial part of the stimulus presentation. It remains the possibility, that although not explicitly encoded, previous magnitude information was maintained in the first part of the stimulus in an ‘activity-silent’ state (Stokes, 2015; Barbosa et al., 2020). Surprisingly, we also discovered that these effects occurred only when the visual features of previous and current stimuli differed, suggesting that the stimulus properties play a triggering role on the influence of prior information.

## Materials and methods

### Results

#### Behavioral results

Two monkeys were trained to discriminate which of two stimuli, S1 and S2, presented sequentially and separated by a first delay (D1) was farther from the center, Fig. 1a.

**Figure 1.**
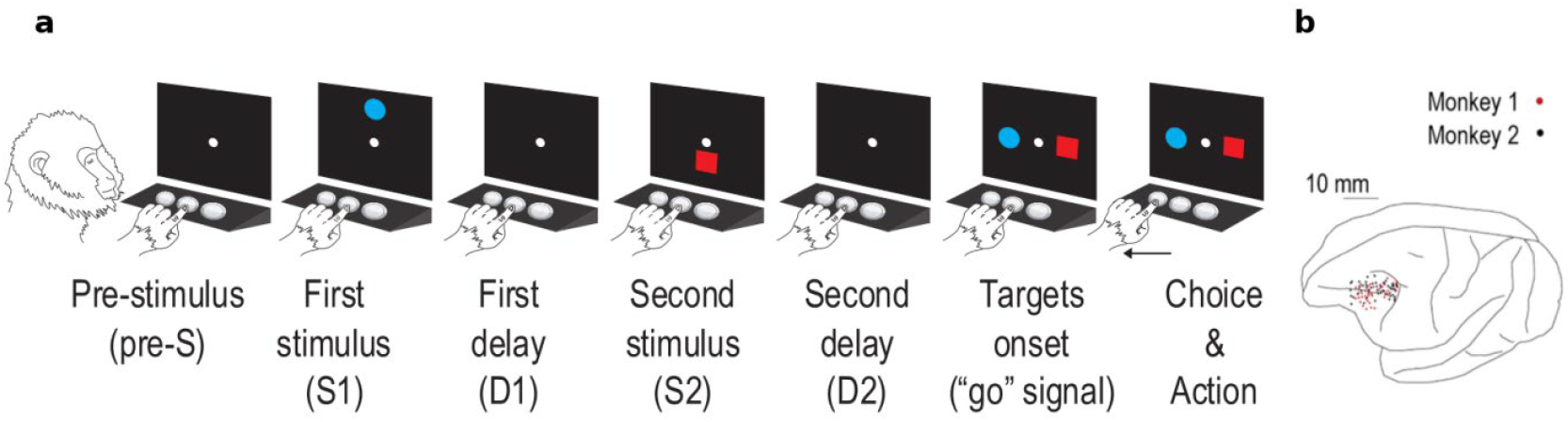
(a) Trial task events. Trials were initiated by the touching of the central switch, which led to the appearance of the central stimulus (reference point). After 400 or 800 ms (pre-stimulus; pre-S), S1 was presented. A variable delay (first delay; D1) of 400 or 800 ms followed S1, after which the second stimulus (S2) appeared. S2 was followed by a second delay (second delay; D2) of 0, 400, or 800 ms. Both S1 and S2 were presented for 1000 ms and presented either above or below the reference point by 8 to 48 mm (8-mm steps). At the target onset, both stimuli reappeared along the horizontal and the monkey had to select the stimulus that had been presented further from the reference point (the blue circle in the example trial) by touching the spatially corresponding switch below the screen. Correct responses were rewarded with 0.1 mL fluid, while errors were followed by acoustic feedback. The stimulus feature (blue circle/red square), position (above/below the reference point), distance from the reference point, and target position (left/right) were pseudo-randomly selected. (b) Penetration sites. Composite from both monkeys, relative to sulcal landmarks.

We tested whether the most recent stimulus S2 presented modulated the contraction bias effect, thereby influencing performance in the current trial. Fig. 2a shows how the magnitude of the previous S2 affected the spatial discrimination in the current trial. We found that as the magnitude of the previous S2 increased, the S2-ward bias in the current trial decreased (linear fit slope=-1.34, p<0.001). In other words, when the previous S2 had a low magnitude, the S1 stimulus in the current trial tended to be underestimated. This resulted in a higher likelihood of the current response being directed towards the S2 stimulus, and conversely, when the previous S2 had a higher magnitude, there was a greater tendency for the current response to favor the S1 stimulus. This is in line with how the contraction bias affects the performance (see inserts in Fig. 2a and Fig. 2b), wherein the first stimulus S1 is contracted toward the mean stimulus distribution (dashed vertical line, drawn in its true position in panel b). The contraction of S1 alters the perception of the distance between the two stimuli, in some cases augmenting it thus making the task easier (green pairs in Fig. 2b, bias +) while in others reducing it thus increasing the task difficulty (red pairs in Fig. 2b, bias -).

**Figure 2.**
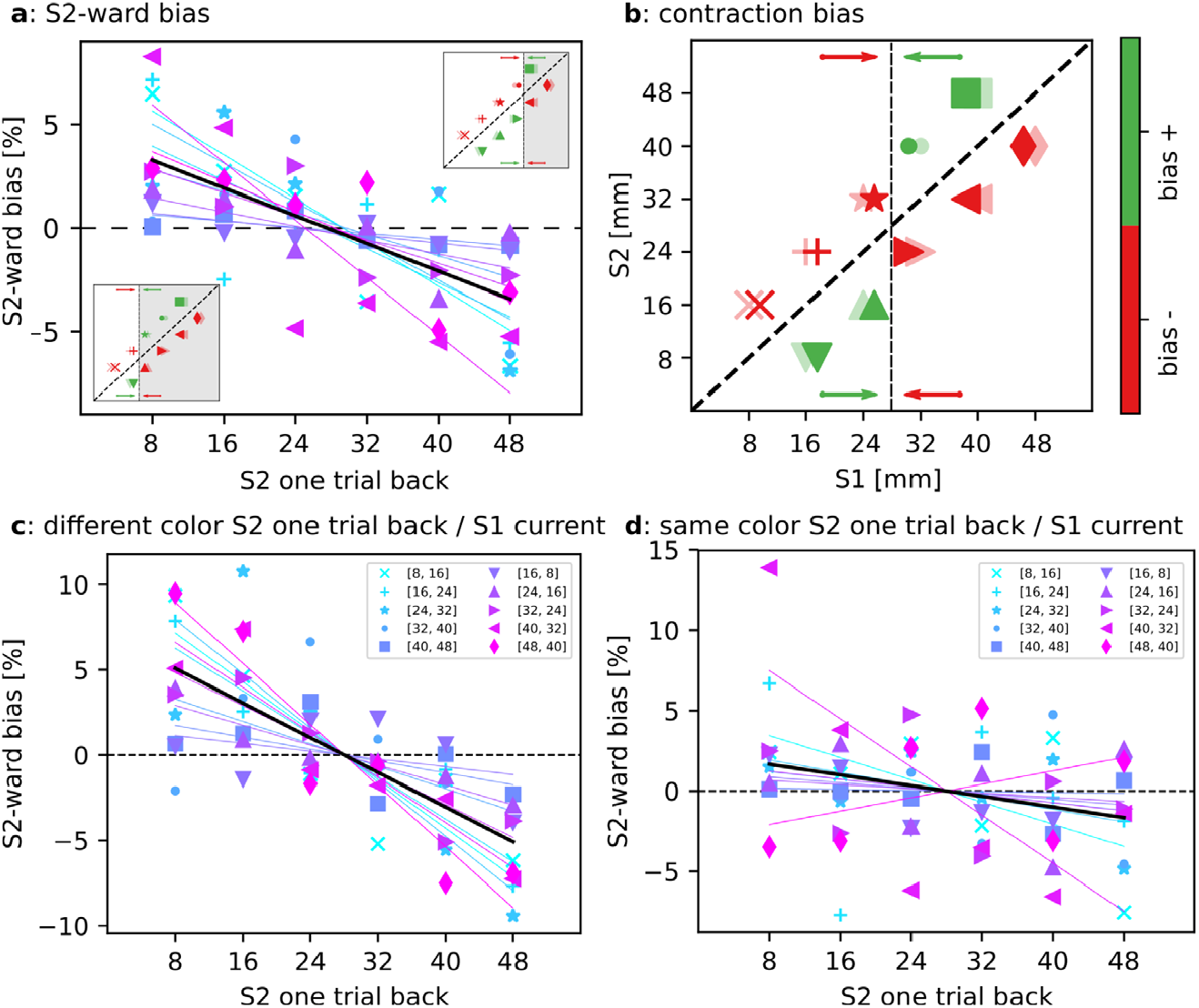
(a) Effect of the previous S2 distance on the performance. For each stimulus pair, the S2-ward bias relative to the previous occurrence of S2 is plotted (colored marker), i.e. the percentage of trials in which S2 was chosen minus the global mean performance of the pair itself. A lower value of the previous S2 tends to elevate the proportion of S2 choices in the present trial, and vice-versa (linear fit slope=-1.34, p=0.001). (b) Assuming an exact stimulus mean perception (black dashed vertical line, 28 mm), the contraction bias manifests in both positive (green) and negative (red) ways in the current trial; previous S2 alters the stimulus mean perception: small values of the previous S2 determine a shift in S1 representation toward lower values that is thus consequently underestimated resulting in a higher percentage of S2 choices (bottom left insert in (a): gray area includes pairs in which S1 is perceived as smaller thus favoring the choice of S2, the opposite in the top right insert: due to a shift toward a higher value of the mean, the gray area becomes smaller, thus the choice of S2 is disfavored). Panels (c) and (d) depict the same concepts as in (a), but with a differentiation based on color mismatch and color match between the previous S2 and the current S1. The influence of the previous S2 persists when there is a color mismatch (linear fit slope=-1.97, p=0.003), while it reduces when colors are matched (linear fit slope=-0.70, p=0.02).

Following this reasoning, the behavioral results show that the recent trial history influenced the perceived mean of the stimulus. A shift of the stimulus mean towards smaller values as a consequence of a small previous S2, determines a higher probability of underestimating S1 and thus a positive S2-ward bias (see bottom left insert in Fig. 2a, the gray area shows where S1 is underestimated), and the opposite for high recent history (see top right insert in Fig. 2a, the smaller gray area indicates a lower probability of underestimating S1). In the lower two panels of Fig. 2, we distinguished trials based on whether there was a color mismatch or a color match between the previous S2 and the current S1.

On average, we observed that the S2-ward bias was more pronounced when there was a color mismatch (linear fit slope=-1.97, p=0.003). Conversely, the S2-ward bias was notably reduced when there was a color match (linear fit slope=-0.70, p=0.02). Here, we only used stimulus pairs with the smallest relative distance, i.e. 8 mm, to avoid other influences on task difficulty. A complete representation of the phenomenon is shown in Fig. S1, which illustrates the effect across all stimulus pairs. Furthermore, to corroborate the relationship between S2-ward bias and the previous S2 stimulus, Fig. S2 reports the effect of the previous S1 stimulus on the current S2-ward bias, showing that it had no effect on the current trial. This comparison is important to exclude repeat/shift response biases.

#### Neural results: decoding of S1 and influence of the previous S2 stimulus

To investigate the influence of the previous trial in terms of S2-ward bias on the neural activity, we focused our analysis on the period of presentation of S1 using the prefrontal recordings of the original dataset. First, we characterized the neural decoding of S1 itself during its presentation. Fig. 3(a-b), shows the decoding of S1 starting from 400 ms before its presentation (0 ms) and continuing for the subsequent 1400 ms (comprising 1000 ms of presentation time and an additional 400 ms delay, D1). The stimulus magnitudes ranged from 8 to 48 mm, with an 8 mm increment, and were categorized into three broader classes: low (8, 16 mm), medium (24, 32 mm), and high (40, 48 mm). As expected, we observed a significant decoding accuracy for S1 both throughout the entire presentation period and during D1 (p<0.05 random permutation test, n=1000; in Fig. 3(a-b) significant results are shown by the gray bar over the time axis, and by the colored area on the map, assuming white indicates non-significance). In particular, panel b shows how the decoding performance changed dynamically when evaluated across all pairs of training and testing time windows (diagonal elements: same training and testing time windows, as shown in panel (a); off-diagonal elements: different training and testing time windows). This dynamic coding indicates that a different coding scheme is used by the prefrontal population of neurons for coding S1 in time. Fig. 3(c-d), similarly to the previous panels, shows the decoding of S1 categorizing the trials into those with a color mismatch (diffColor) and those with a color match (sameColor) between the previous S2 and the current S1. This categorization had a significant impact on the decoding performance, leading to a notable decrease, likely caused by the reduction to half of the number of trials and the disentanglement of the color matching feature. However, it is worth noting that there still remains a higher decoding accuracy in the diffColor condition, which aligns with the behavioral data as depicted in Fig. 2(c-d).

**Figure 3.**
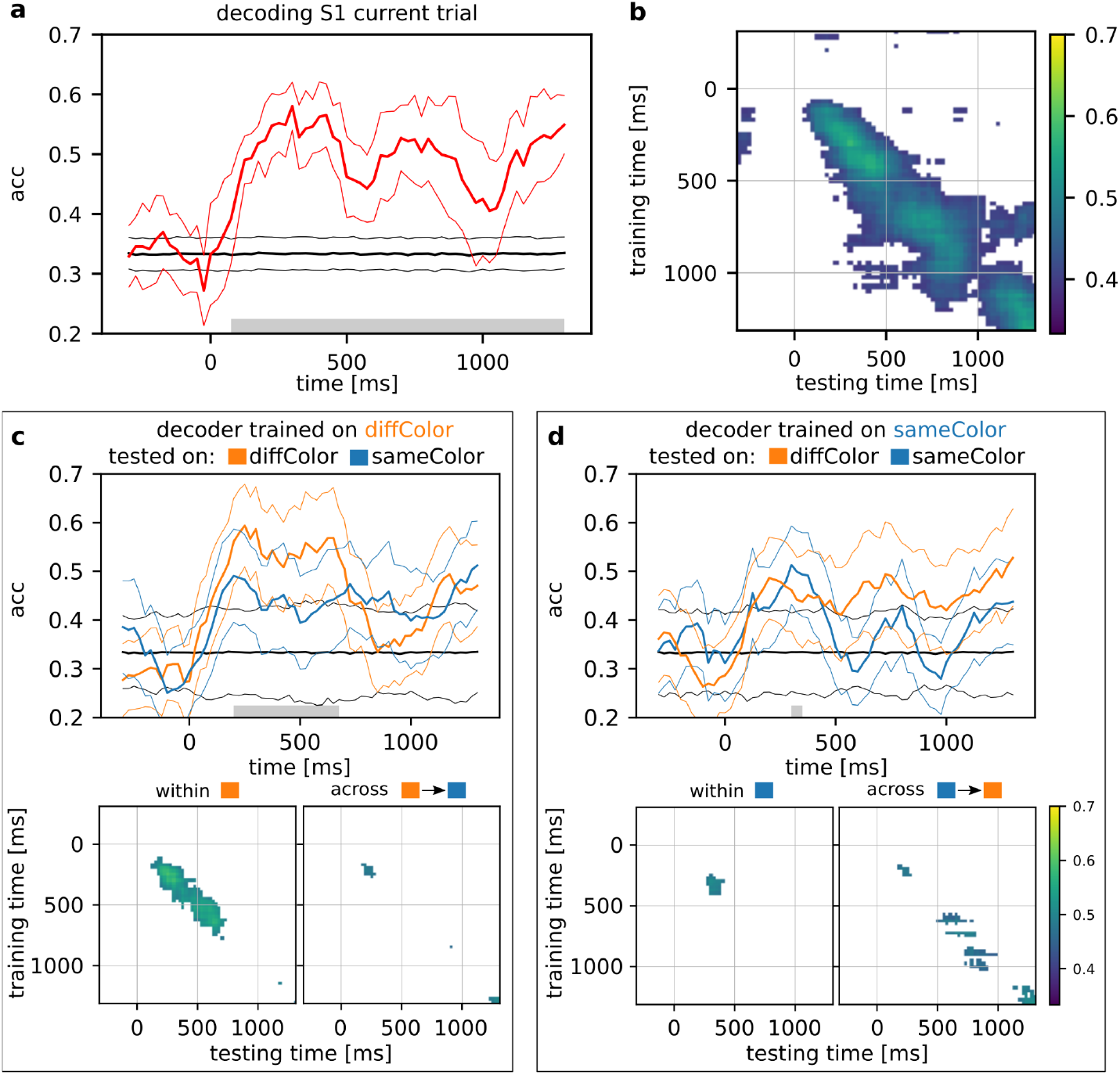
(a) Decoding accuracy of stimulus S1 ± s.d. grouped in three macro classes: low (8, 16 mm), medium (24, 32 mm) and high (40, 48 mm). Accuracy is shown from 400 ms before S1 appearance (0 ms) until 400 ms after the end of its presentation period (which lasted for 1000 ms, the final 400 ms is the shortest delay D1 in common with all trials) (bin size: 200 ms, step size: 25 ms, 100 repetitions with random train/test split). The gray bar over the time axis indicates a significant difference from chance (random permutation test, 1000 iterations). (b) Same decoding performance evaluated across training and testing time window (the diagonal corresponds to the curve in panel (a)). Panels (c) and (d) illustrate the same concept as depicted in panels (a,b), but with a distinction based on whether there is a color mismatch (diffColor) or color match (sameColor) between the previous S2 and the current S1. Decoding accuracies for both within- and across-condition analyses are displayed, accompanied by corresponding training/testing time maps (bottom section). White shading in the maps indicates no significant deviation from chance levels.

To assess the impact of the previous trial on the decoding accuracy of S1, we conditioned its classification on the S2 distance of the preceding trial. We maintained the same macro-level groups (low, medium, and high magnitude) for the previous S2 as well and clustered each trial accordingly. Given the randomization of stimuli across trials, we anticipated a balanced distribution of the S1 classes (low, medium, and high) within each S2 one-trial back class. This approach enabled us to characterize the neural decoding of S1 for each category of S2 one-trial back. Fig. 4(a-d), show the mean decoding accuracy of S1 within and across the previous S2 conditions. Specifically, panels (a, c) refer to the first (early S1) and second (late S1) half periods of S1 presentation, respectively.

**Figure 4.**
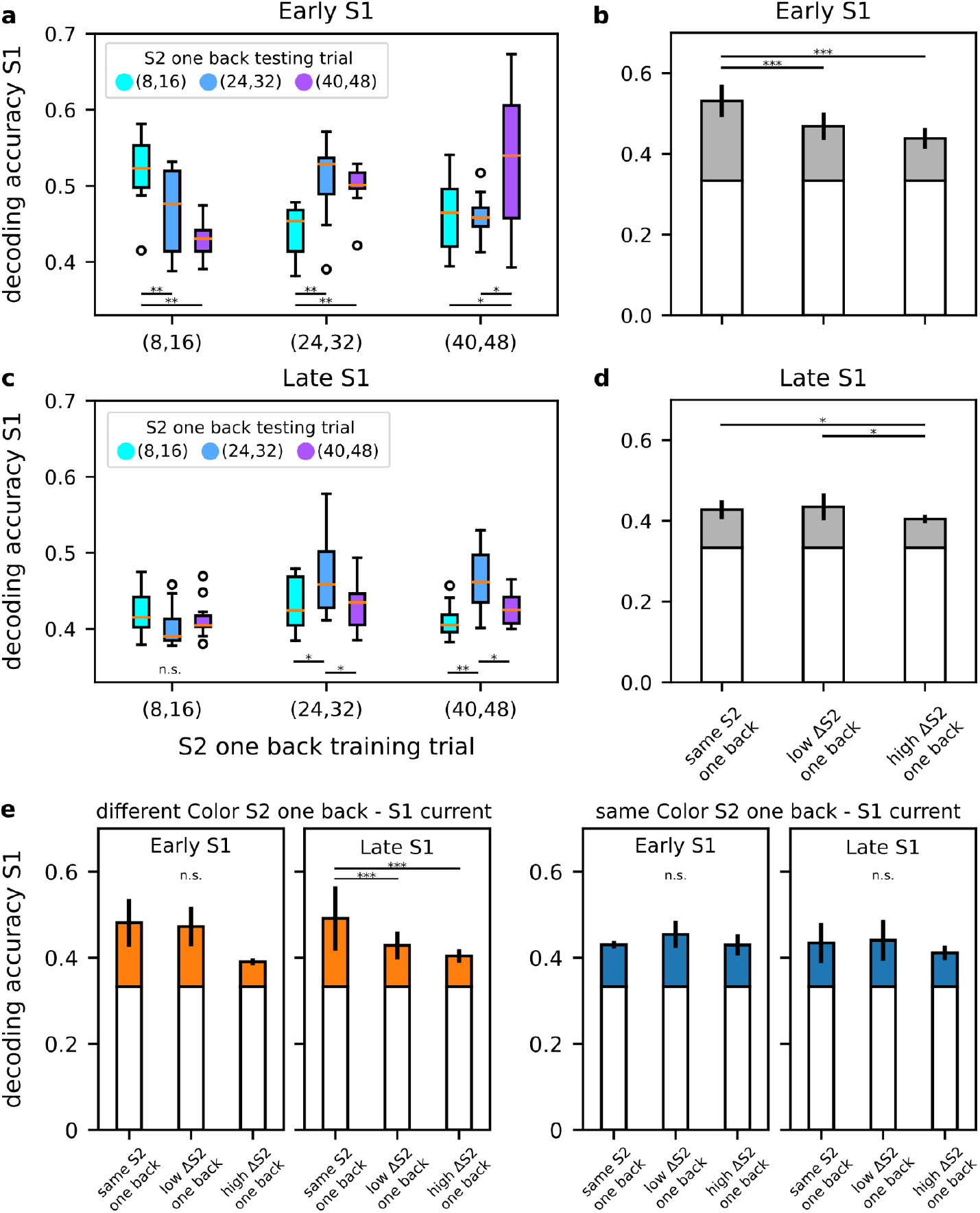
(a) Decoding accuracy of stimulus S1 conditioned on S2 of the previous trial, boxplots show how the accuracy changes in the first half of S1 presentation (early S1) when training and testing trials share (or not) their S2 in one-trial back. (b) Same results as in (a) but grouped in: decoding of S1 within condition (same previous S2 trial) and across conditions (low ΔS2: small gap between S2 one-trial back in train and test sets, e.g. (8,16) vs. (24,32), high ΔS2: large gap between S2 one-trial back in train and test sets, e.g. (8,16) vs. (40,48)). Similarly, in panels (c-d), focusing on the last half of the S1 presentation (late S1). In panel (e), following a structure analogous to that of panels (b) and (d), but with trials divided for color mismatch (orange bars) or match (blue bars) between the previous S2 and the current S1. n.s.=not significant, *=p<0.05, **=p<0.01, ***=p<0.001, ANOVA test with Tukey’s multiple comparison test.

In general, we observed that the highest decoding accuracy was achieved when the training and testing trials used to decode S1 had similar previous S2 values, indicating an advantage of decoding S1 within the same S2 one-trial back condition. These accuracy values closely align with the results in Fig. 3a, where the previous trial’s S2 was not taken into account. However, the ability to decode S1 significantly diminished in cases where there was a discordant S2 one-trial back condition. Notably, the reduction in accuracy corresponded to the degree of disparity with the S2 values from one-trial back: the greater the discrepancy, the more pronounced the decline in S1 accuracy. This pattern is visually represented in Fig. 4(b,d). The column labeled “same S2 one back” pertains to within-condition decoding, while the other two columns represent across-condition decoding, which was further divided into “low ΔS2 one back” and “high ΔS2 one back”. “Low ΔS2 one back” indicates low discrepancy between S2 one-trial back in training and testing trials, e.g. *low* S2 one back in training and *medium* in testing, on the contrary “high ΔS2 one back” indicates high discrepancy, e.g. *low* S2 one back in training and *high* in testing. This result was particularly prominent in the early S1 period (as seen in panel b) and gradually diminished as S1 progressed into its later stages (as observed in panel d), eventually fading during the D1 period, as shown in Fig. S3.

In summary, these findings indicate that when a decoder, initially trained to distinguish S1 using trials with a specific S2 history, is subsequently tested in a trial with a different S2 history, its performance declines. This result suggests that the S2 stimulus from one-trial back has influenced the perception of S1 in the current trial, leading to a disruption in the decoder’s ability to accurately classify S1.

In the lower part of Fig. 4, panel (e) presents the results of the same analysis, which was conducted separately for the diffColor and sameColor conditions. Consistent with the previous observations, the diffColor condition mirrored the overall results obtained without trial separations, with the difference that a significant effect was found in the late S1, while only a tendency was maintained in early S1. However, in the sameColor condition, the previously observed effect disappears.

#### Neural results: HMM analysis and population decoding of S1 color

The preceding section provided evidence of interference from the previous S2 on the decoding of the current S1, indicating that the stimulus features can influence this process. This suggests the possibility that both variables, namely S2 from one-trial back and S1 color, might be decoded during the presentation of S1, as they both contributed to determining the ability to decode the magnitude of S1.

Regarding the color feature, we studied each recording session separately by applying a hidden Markov model (HMM) analysis and also through a population decoding approach similar to what was described in the previous section. The HMM analysis aims to model the dynamics of neural activity in an ensemble of simultaneously recorded neurons. This approach is based on the assumption that neural dynamics evolve as a sequence of hidden states, each characterized by a specific firing profile. Thus, it aims to segment neural activity into sequences of alternating metastable states. Interestingly, sequences of HMM states have already been shown to reliably characterize this dataset (Benozzo et al., 2021). Specifically, coding states have been reported during the presentation of S2, coding for the relative distance between the two stimuli based on their color, i.e. S_red_ greater or lower than S_blue_, and their order of presentation, i.e. S1 greater or lower than S2. Here, unlike our previous work, we extended the time window for each trial to include the initial phase of the next trial. This extension allowed us to capture a sequence of states encompassing the distance discrimination process, the subsequent response, and related feedback up to the presentation of stimulus S1 in the next trial. This enabled us to study state dynamics during S1, giving the hidden state the ability to potentially capture traces from the previous trial. Testing for coding states of S1 color separately in diffColor and sameColor trials yielded the following results: 6 out of 41 sessions contain a state with mean occupancy that significantly differs between red and blue stimuli in the diffColor condition, while 5 out of 41 sessions do so in the sameColor condition. Interestingly, all these coding states are condition specific, meaning that if a state codes for color in one condition, it does not do so in the other (with the exception of one session where the same state codes for both conditions). Fig. 5a shows two examples of sessions with coding states for the stimulus color feature. The panel reports the mean state occupancy for each state, separately for the two conditions and the two colors, computed both during the stimulus presentation window (first row) and during the pre-stimulus delay (second row). Subplots with reduced transparency refer to states that are not coding states, highlighting that none of them are coding states during the pre-stimulus delay. The last row of the panel shows the firing rate profile for each state. A more comprehensive visualization of state sequences is shown in Fig. S4, where the HMM state sequence for each coding state is reported at the single trial level, spanning from the appearance of S2 in the previous trial to the end of S1 presentation in the current trial.

**Figure 5.**
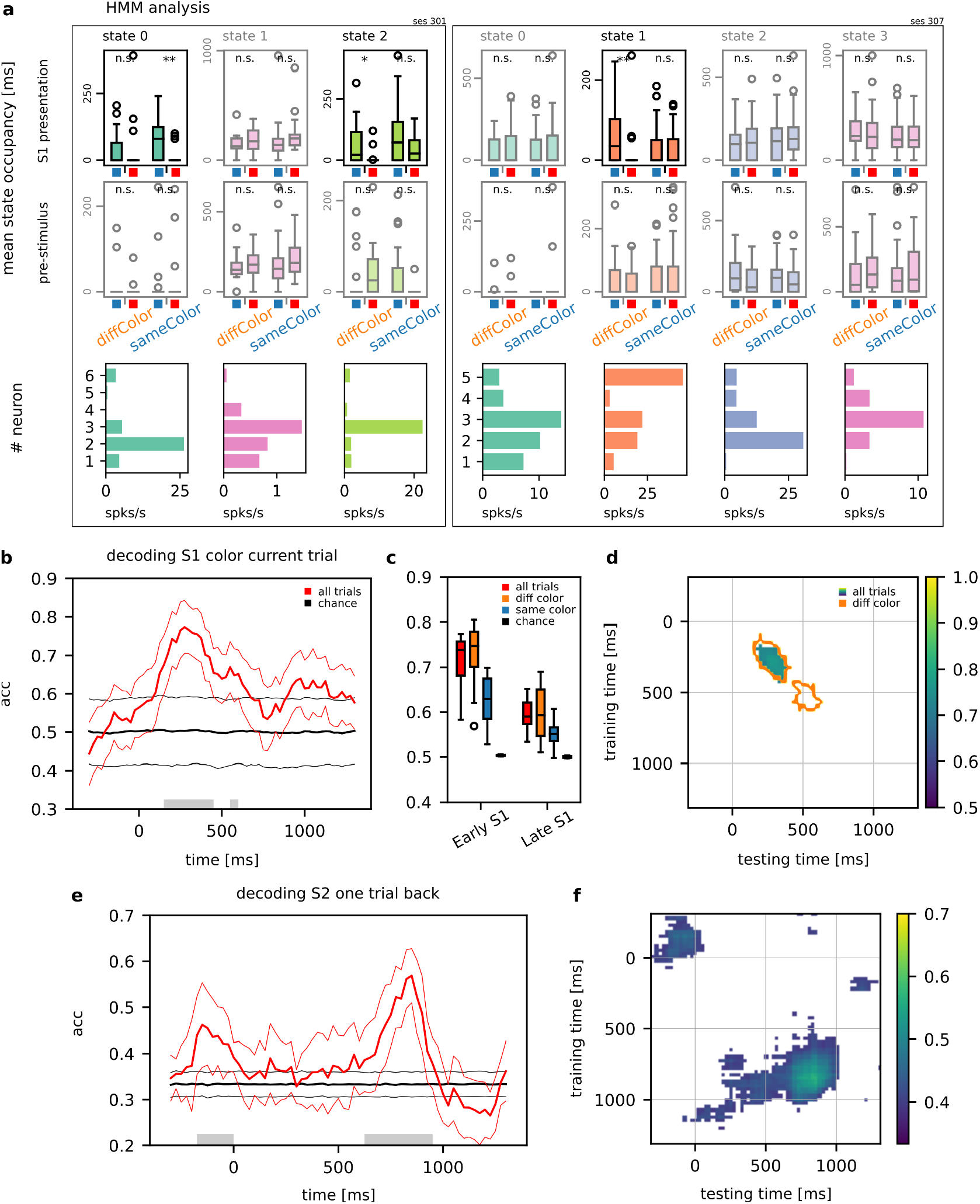
(a) Examples of sessions with significant HMM coding states for the first stimulus color feature. In each session’s panel, the mean state occupancy is depicted for each state, condition (diffColor vs. sameColor) and stimulus color (blue vs. red). The first row refers to the time window of S1 presentation, while the second row covers the pre-stimulus delay interval. States with no significant mean occupancy between colors are displayed with reduced transparency (n.s.=not significant, *=p<0.05, **=p<0.01, Mann-Whitney U rank test with Benjamini/Hochberg false discovery rate correction). The last row shows the firing rate profile of each state. (b) Decoding accuracy of the color (blue vs. red) of stimulus S1 ± s.d. Accuracy is shown from 400 ms before S1 appearance (0 ms) until 400 ms after the end of its presentation period (which lasted for 1000 ms, the final 400 ms is the shortest delay D1 in common with all trials) (bin size: 200 ms, step size: 25 ms, 100 repetitions with random train/test split). The gray bar over the time axis indicates a significant difference from chance (random permutation test, 1000 iterations). In panel (c), boxplots compare decoding accuracies when S1 color mismatches (diffColor, orange boxplot) or matches (sameColot, blue boxplot) with the color of the previous S2; the red boxplot reports the global accuracy without separating trials on color conditions. (d) Decoding performances evaluated across different training and testing time windows: the map illustrates the scenario without trial separation, the contour lines delineate the expansion of the map in the context of color mismatch, while no significant accuracies were observed when colors matched. (e) Decoding accuracy of the previous S2 ± s.d. during the S1 presentation (current trial). The previous S2 stimulus distances are grouped in three macro-classes: low, medium and high distances. Same approach as in panel (b). (f) The decoding performance evaluated across different training and testing time windows.

Regarding the population decoding analysis, we also assessed the ability to classify each trial based on the color of S1. In Fig. 5b, the accuracy curve is presented, revealing a significant peak during the first half of the presentation window. In panel (c), the mean accuracy values for “early S1” and “late S1” are reported (red boxplots). Additionally, the panel includes the mean accuracy values for these two periods in both the diffColor and sameColor conditions, illustrated in orange and blue boxplots, respectively. In line with our previous findings, the diffColor condition enhanced the discriminability of the stimulus features. In panel (d), is shown the training/testing time map. This map illustrates the scenario without trial separation, with contour lines delineating the expansion of the map in the context of color mismatch (diffColor), while no significant accuracies were observed when the colors matched (sameColor). Both analyses provided evidence of the importance of the visual features in the neural data, specifically showing a relationship with the color feature of the previous stimulus.

Focusing on S2 of one-trial back, despite its irrelevance to task performance but due to its impact on S1 decoding, we explored whether there was still a trace of it in the current trial. Fig. 5e illustrates that after an initial encoding of the magnitude of the previous S2 (before the presentation of S1, 0 ms), it was no longer decoded until the second half of S1 presentation (late S1). During this last period, it reached its highest accuracy, nearly 0.6, while it was at chance level in the early S1 period. Interestingly, this peak of accuracy for decoding the previous S2 appeared later than the time window in which the effect of S2 history on the decoding of S1 was observed. This raises the possibility that a silent trace of the previous S2 distance might influence the coding of S1 without being explicitly represented, as we will propose in the Discussion. In panel (f), the decoding performance is visualized across all pairs of training and testing time windows. Two distinct clusters are evident along the diagonal. The first, with lower intensity, is located before 0 ms, corresponding to the initial peak observed in panel (e). The second cluster, the most prominent one, is centered in the late S1 period, aligning with the highest peak in panel (e). The white area in the map signifies no significant decoding during those time intervals (random permutation test n=1000, p<0.01). This panel suggests that there was no discernible relationship between the coding of S2 before S1 and the late S1 period, as no significant decoding was observed across these two periods. This lack of significance hints at a potential change in the population coding between these time intervals, indicating that the processing of information changed at the population level across different neurons as the trial progressed from the early to late S1 period. As we did for the color matching feature in the study of S1 magnitude decoding, we also examined the effect of color match/mismatch when applied to S2 one-trial back. However, no significant results were observed in either the match or mismatch conditions (not shown).

## Discussion

We studied the influence of the most recently presented stimulus on the performance of monkeys in a distance discrimination task to investigate its neural bases in the prefrontal cortex and whether we can find signatures already in the stimulus coding of this influence. Specifically, we investigated whether the most recent stimulus could generate a contraction bias effect and interfere with the coding of the stimulus to discriminate. Consistent with the findings of Benozzo et al. (2023) regarding the average distribution of stimuli, we also found short term effects of the previous trial with the previous S2 significantly affecting the performance, contributing to the modulation of the contraction bias effect. Specifically, we identified a contraction bias effect of the previous S2 magnitude which appeared as a shift in the representation of the S1 magnitude toward the previous S2. As a possible neural substrate, we found that the magnitude of the previous S2 affected the decoding of S1. S1 was decoded with the highest accuracy when the decoder was trained and tested with S2 of the same magnitude, while conversely, a mismatch in magnitudes altered its decoding, as shown by the reduction in the decoding. The greater the mismatch, the higher the reduction in accuracy. As discussed in a later section, we also found that the effect of the previous trial depended on whether the visual features of S1 differed from those of the previously presented S2.

The effect of the previous S2 on the representation of S1 suggests that biases generated from the previous trial can affect the representation of the stimulus to be discriminated and not only the decision process, as we have already shown in our previous work (Benozzo et al., 2023). In that study, we demonstrated that the decision process is affected by the mean distribution of stimulus magnitudes, but we were unable to determine whether the representation of the first stimulus’s magnitude was also influenced by contraction bias. This limitation arose because comparing the effect of different mean distributions on the encoding of the first stimulus would have been necessary. In the present study, we address this limitation by investigating how the previous trial affects the encoding of the first stimulus when the second stimulus has varying magnitudes. Our findings reveal that biases from previous trials can indeed alter the representation of the stimulus to be discriminated. This suggests that the decision-related activity observed after both stimuli are presented may be driven, at least in part, by a biased representation of the first stimulus’s magnitude at the time it is presented.

Our results for the influence of previous trials, remind the findings by Histed et al., (2009) on the effect of history on the current trial coding activity. They showed that the previous outcome information was maintained across trials and that task variables’ performance and coding, such as response direction, were encoded stronger after correct than error trials, possibly favoring learning. Our result is also in line with the behavioral results of Papadimitriou et al., (2015) in monkeys, who found that memory in the current trial shows a bias toward the location of the memorandum from the previous trial (proactive interference). The effect of the history bias on performance was also reported in rats by Akrami et al., (2018), where the activity of the posterior parietal cortex revealed a causal role in modulating the impact of recent trials on performance. Similarly, Barbosa et al., (2020) found a stronger effect of previous trials linked to a reactivation of activity in the monkey and human prefrontal cortex during the initial phase of the trial, which was previously silent during the inter-trial interval. More specifically, regarding the role of the previous S2 in affecting the decoding of the current S1 (see Fig. 4), Boboeva et al., (2024) recently developed a computation model that aims to reproduce the interplay between the parietal cortex and the working memory network. Their model showed a direct relationship between the contraction of S1 and its difference from the previous trial’s stimulus S2. This indeed strongly resonates with our findings, where S1 decoding is maximized when the decoder was trained and tested on trials with similar previous S2 magnitude, and proportionally decreased with increasing their discrepancy.

We searched for a trace of the past S2 magnitude during the S1 presentation that could account for the previous trial effects. We found that the previous S2 modulated the current trial activity in the pre-stimulus period but decreased to chance levels after the presentation of S1. Its modulation reappeared again in the latest S1 period, as if the memory of the previous S2 resurfaced only after a period of no explicit representation. The first study showing an interruption of coding in time, conceptualized as a silent activity, is a study by Watanabe and Funahashi (2014). Recording from the prefrontal cortex, they found that using a dual task paradigm, the memory activity associated with a location to be held in memory disappeared temporarily during an intervening attentional task, only to emerge again later when the concurrent task was completed. The reactivation of the representation of the previous stimulus in the pre-stimulus interval right before the appearance of the new stimulus also resembles the effect observed by Barbosa et al. (2020) and represents a confirmation of this phenomenon. According to Barbosa and colleagues, a break in the continuity of coding should be interpreted not as a loss of the information in the network but as a sign of a switch to a different modality of information maintenance, termed ‘activity-silent’. Being in an ‘activity-silent’ state has been proposed to support working memory (Stokes, 2015). The mechanism underlying ‘activity-silent’ states is not yet clear; one proposed mechanism is that information is maintained in memory through a temporary modification of the synaptic weights (Mongillo et al., 2008; Barak et al., 2014; Kaminsky et al., 2020). ‘Activity-silent’ network dynamics would represent an alternative mechanism to persistent and dynamic coding (Meyers et al., 2008; Meyers et al., 2012; Mendoza-Halliday and Martinez-Trujillo, 2017; Meyers et al., 2018; Ceccarelli et al., 2023) for keeping information in working memory. In addition to neurophysiology, some evidence for activity-silent forms of memory maintenance comes from neuroimaging studies e.g., using retro-cue paradigms (Rose et al., 2016). However, imaging studies have the limitation that they cannot determine whether the absence of decoding capability reflects a complete loss of coding activity at the single cell level or just a reduction of the cells involved (Kaminsky et al., 2020).

Why information is kept in an ‘activity-silent’ state in some cases but not in others remains unclear. In our study, the reactivation of the previous S2 representation aligned temporally in the late S1 with the period in which it affected mostly the decoding of S1, which is in trials when the visual stimulus features differed between S1 and the previous S2. It is not clear why an explicit representation of S2 was lost during the initial representation of S1. One possible interpretation is that the shift to an ‘activity-silent’ representation resulted from the interaction between current and past neural representations, where neural resources are prioritized for processing the current stimulus over the past one because it is more relevant. It is possible that the resources allocated in our task for other computations, such as the representation of the stimulus features of the first stimulus, dominated the competition for relevance over the information about the previous stimulus.

We also observed an unexpected effect: the influence of matching of visual stimulus features across trials on the emergence of the bias. Why does the bias of the last trial on behavior go away when the stimulus presented shares the same visual features as the previous S2? One tentative explanation is that the reduction of the first stimulus feature coding depends on an adaptation effect (Gross et al., 1967; Barron et al., 2016; Henson, 2003; Karlaftis et al., 2021). This effect has been shown also in monkeys, not only in the IT cortex (Gross et al., 1967; Miller et al., 1991) but also in the prefrontal cortex by Miller et al., (1996) and Rainer et al., (1999). Karlafti and colleagues (2021) have shown in an fMRI study in humans a decrease in activity in the dorsolateral prefrontal cortex following the repeated presentation of sensory stimuli. We hypothesize that the effect of the repetition of the same visual features affects other computations. First, it would reduce the response to the visual features of the first stimulus presentation. Second, the adaptation could simultaneously influence concurrent representations, such as the magnitude of S1 (Fig. 3d) and the impact of information from the previous S2 on S1 coding (Fig. 4e, right panel). This hypothesis is further supported by the computational model of the working memory network proposed by Boboeva et al., (2024), which includes an adaptation term that adjusts the persistence of self-sustained activity related to past stimuli. The stronger the adaptation term, the faster the extinguishing process of the previous trial’s activity. Currently, this remains speculative, and future studies are needed to understand this effect and to test whether it can be generalized to humans and other species and to other paradigms and sensory modalities.

## Acknowledgements

We thank Dr Satoshi Tsujimoto, Dr Steven P. Wise for their contribution to the experimental and data acquisition aspects and Dr Andrew Mitz for engineering.

## Funding

This work was supported by by Ministero dell’Università e della Ricerca (Grant PRIN 2017: 2017KZNZLN_004).

## Notes

### Competing Interest Statement

The authors have declared no competing interest.

## References

Akrami A, Kopec CD, Diamond ME, Brody CD. Posterior parietal cortex represents sensory history and mediates its effects on behaviour. Nature 554, 368–372 (2018).

Ashourian P, Loewenstein Y. Bayesian inference underlies the contraction bias in delayed comparison tasks. PLoS ONE 6(5), e19551 (2011).

Barak O, Tsodyks M. Working models of working memory. Curr Opin Neurobiol. 25, 20–4 (2014).

Barbosa J, Stein H, Martinez RL, Galan-Gadea A, Li S, Dalmau J, Adam KCS, Valls-Solé J, Constantinidis C, Compte A. Interplay between persistent activity and activity-silent dynamics in the prefrontal cortex underlies serial biases in working memory. Nat Neurosci. 23(8), 1016–1024 (2020).

Barraclough DJ, Conroy ML, Lee D. Prefrontal cortex and decision making in a mixed-strategy game. Nat Neurosci. 7(4), 404–410 (2004).

Barron HC, Garvert MM, Behrens TE. Repetition suppression: a means to index neural representations using BOLD? Philos Trans R Soc Lond B Biol Sci. 5, 371(1705):20150355 (2016).

Bausenhart KM, Bratzke D, Ulrich R. Formation and representation of temporal reference information. Curr Opin Behav Sci. 8, 46–52 (2016).

Benozzo D, Ferrucci L, Genovesio A. Effects of contraction bias on the decision process in the macaque prefrontal cortex. Cereb Cortex. 10, 33(6):2958–2968 (2023).

Benozzo D, La Camera G, Genovesio A. Slower prefrontal metastable dynamics during deliberation predicts error trials in a distance discrimination task. Cell Rep. 6, 35(1):108934 (2021).

Boboeva V, Pezzotta A, Clopath C, Akrami A. Unifying network model links recency and central tendency biases in working memory. eLife 12, RP86725 (2024).

Ceccarelli F, Ferrucci L, Londei F, Ramawat S, Brunamonti E, Genovesio A. Static and dynamic coding in distinct cell types during associative learning in the prefrontal cortex. Nat Commun. 14, 14(1):8325 (2023).

Cicchini GM, Arrighi R, Cecchetti L, Giusti M, Burr DC. Optimal encoding of interval timing in expert percussionists. J Neurosci. 32(3), 1056–1060 (2012).

Fassihi A, Akrami A, Pulecchi F, Schonfelder V, Diamond ME. Transformation of Perception from Sensory to Motor Cortex. Curr. Biol. 27, 1585–1596 (2017).

Genovesio A, Brasted PJ, Mitz AR, Wise SP. Prefrontal cortex activity related to abstract response strategies. Neuron. 47(2), 307–20 (2005).

Genovesio A, Tsujimoto S, Wise SP. Prefrontal cortex activity during the discrimination of relative distance. J Neurosci. 31(11), 3968–80 (2011).

Genovesio A, Ferraina S. The influence of recent decisions on future goal selection. Philos Trans R Soc Lond B Biol Sci. 369(1655), 20130477 (2014).

Genovesio A, Cirillo R, Tsujimoto S, Mohammad Abdellatif S, Wise SP. Automatic comparison of stimulus durations in the primate prefrontal cortex: the neural basis of across-task interference. J Neurophysiol. 114(1), 48–56 (2015).

Genovesio A, Seitz LK, Tsujimoto S, Wise SP. Context-Dependent Duration Signals in the Primate Prefrontal Cortex. Cereb Cortex. 26(8), 3345–56 (2016).

Gross CG, Schiller PH, Wells C, Gerstein GL. Single-unit activity in temporal association cortex of the monkey. J Neurophysiol. 30(4), 833–43 (1967).

Henson RN. Neuroimaging studies of priming. Prog Neurobiol. 70(1), 53–81 (2003).

Hermoso-Mendizabal, A., Hyafil, A., Rueda-Orozco, P.E. et al. Response outcomes gate the impact of expectations on perceptual decisions. Nat Commun. 11, 1057 (2020).

Histed MH, Pasupathy A, Miller EK. Learning substrates in the primate prefrontal cortex and striatum: sustained activity related to successful actions. Neuron. 63(2), 244–53 (2009).

Hollingworth HL. The central tendency of judgment. J Philos Psychol Sci Method. 7(17), 461–469 (1910).

Kamiński J, Rutishauser U. Between persistently active and activity-silent frameworks: novel vistas on the cellular basis of working memory. Ann N Y Acad Sci. 1464(1), 64–75 (2020).

Karlaftis VM, Giorgio J, Zamboni E, Frangou P, Rideaux R, Ziminski JJ, Kourtzi Z. Functional Interactions between Sensory and Memory Networks for Adaptive Behavior. Cereb Cortex. 31(12), 5319–5330 (2021).

Jazayeri M, Shadlen MN. Temporal context calibrates interval timing. Nat. Neurosci. 13, 1020–1026 (2010).

Maboudi K, Ackermann E, de Jong LW, Pfeiffer BE, Foster D, Diba K, Kemere C. Uncovering temporal structure in hippocampal output patterns. Elife. 7, e34467 (2018).

Marcos E, Tsujimoto S, Genovesio A. Independent coding of absolute duration and distance magnitudes in the prefrontal cortex. J Neurophysiol. 117(1), 195–203 (2017).

Marcos E, Tsujimoto S, Mattia M, Genovesio A. A Network Activity Reconfiguration Underlies the Transition from Goal to Action. Cell Rep. 27(10), 2909–2920 (2019).

Mazzucato, L., La Camera, G., Fontanini, A. Expectation-induced modulation of metastable activity underlies faster coding of sensory stimuli. Nat Neurosci. 22, 787–796 (2019).

Mochol G, Kiani R, Moreno-Bote R. Prefrontal cortex represents heuristics that shape choice bias and its integration into future behavior. Curr. Biol. 31, 1234–1244 (2021).

Mendoza-Halliday D, Martinez-Trujillo JC. Neuronal population coding of perceived and memorized visual features in the lateral prefrontal cortex. Nat Commun. 8, 15471 (2017)

Meyers EM. Dynamic population coding and its relationship to working memory. J Neurophysiol. 120(5), 2260–2268 (2018).

Meyers, E.M., Freedman, D.J., Kreiman, G., Miller, E.K., Poggio, T. Dynamic Population Coding of Category Information in Inferior Temporal and Prefrontal Cortex. Journal of Neurophysiology 100, 1407–1419 (2008).

Meyers EM, Qi XL, Constantinidis C. Incorporation of new information into prefrontal cortical activity after learning working memory tasks. Proc Natl Acad Sci U S A. 109(12), 4651–6 (2012).

Miller EK, Li L, Desimone R. A neural mechanism for working and recognition memory in inferior temporal cortex. Science 254(5036), 1377–9 (1991).

Miller EK, Erickson CA, Desimone R. Neural mechanisms of visual working memory in prefrontal cortex of the macaque. J Neurosci. 16(16), 5154–67 (1996).

Mongillo G, Barak O, Tsodyks M. Synaptic theory of working memory. Science 319(5869), 1543–6 (2008).

Olkkonen M, McCarthy PF, Allred SR. The central tendency bias in color perception: Effects of internal and external noise. J. Vis. 14, 5 (2014).

Papadimitriou C, Ferdoash A, Snyder LH. Ghosts in the machine: memory interference from the previous trial. J Neurophysiol. 113(2), 567–77 (2015).

Padoa-Schioppa C. Neuronal origins of choice variability in economic decisions. Neuron 80, 1322–1336 (2013).

Petzschner FH, Glasauer S, Stephan KE. A Bayesian perspective on magnitude estimation. Trends Cogn Sci. 19(5), 285–293 (2015).

Ponce-Alvarez A, Nácher V, Luna R, Riehle A, Romo R. Dynamics of cortical neuronal ensembles transit from decision making to storage for later report. J Neurosci. 32(35), 11956–69 (2012).

Rainer G, Rao SC, Miller EK. Prospective coding for objects in primate prefrontal cortex. J Neurosci. 19(13), 5493–505 (1999).

Raviv O, Ahissar M, Loewenstein Y. How recent history affects perception: the normative approach and its heuristic approximation. PLoS Comput Biol. 8(10), e1002731 (2012).

Raviv O, Lieder I, Loewenstein Y, Ahissar M. Contradictory behavioral biases result from the influence of past stimuli on perception. PLoS Comput Biol. 10(12), e1003948 (2014).

Rigotti M, Barak O, Warden MR, Wang XJ, Daw ND, Miller EK, Fusi S. The importance of mixed selectivity in complex cognitive tasks. Nature 497(7451), 585–590 (2013).

Roach NW, McGraw PV, Whitaker DJ, Heron J. Generalization of prior information for rapid Bayesian time estimation. Proc. Natl. Acad. Sci. U.S.A. 114, 412–417 (2017).

Rose NS, LaRocque JJ, Riggall AC, Gosseries O, Starrett MJ, Meyering EE, Postle BR. Reactivation of latent working memories with transcranial magnetic stimulation. Science 354(6316), 1136–1139 (2016).

Seo, H., Barraclough, D.J., and Lee, D. Dynamic signals related to choices and outcomes in the dorsolateral prefrontal cortex. Cereb. Cortex 17 (Suppl 1), i110–i117 (2007).

Seo, H., and Lee, D. Behavioral and neural changes after gains and losses of conditioned reinforcers. J. Neurosci. 29, 3627–3641 (2009).

Shi Z, Burr D. Predictive coding of multisensory timing. Curr. Opin. Behav Sci. 8, 200–206 (2016).

Stokes MG. ‘Activity-silent’ working memory in prefrontal cortex: a dynamic coding framework. Trends Cogn Sci. 19(7), 394–405 (2015).

Tsujimoto S, Postle BR. The prefrontal cortex and oculomotor delayed response: a reconsideration of the “mnemonic scotoma”. J Cogn Neurosci. 24(3), 627–35 (2012).

Watanabe K, Funahashi S. Neural mechanisms of dual-task interference and cognitive capacity limitation in the prefrontal cortex. Nat Neurosci. 17(4), 601–11 (2014).

Wirth, S., Avsar, E., Chiu, C.C., Sharma, V., Smith, A.C., Brown, E., and Suzuki, W.A. Trial outcome and associative learning signals in the monkey hippocampus. Neuron 61, 930–940 (2009).

Zhang H, Zhou X.. Supramodal representation of temporal priors calibrates interval timing. J. Neurophysiol. 118:2, 1244–1256 (2017).

