## Supplementary for "History bias and its perturbation of the stimulus representation in the macaque prefrontal cortex"

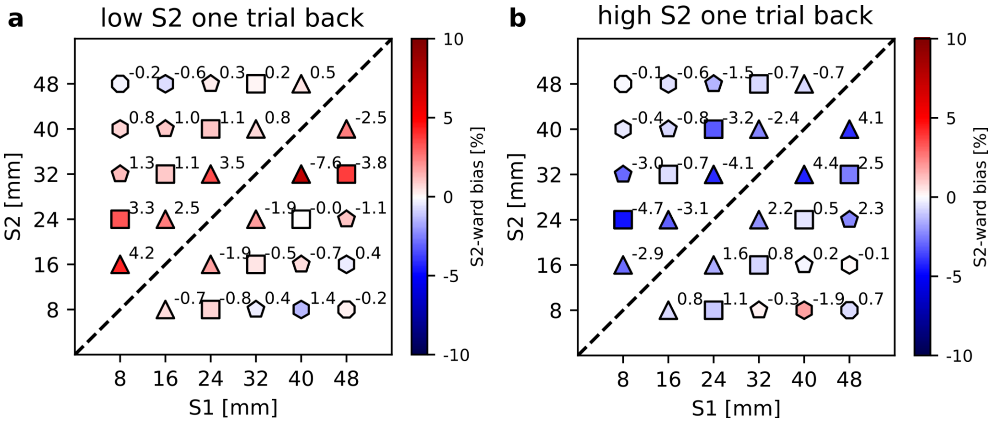


Figure S1. Effect of the previous S2 on the performance of the current trial showed for all stimulus pairs by dividing trials based on the value of the previous S2. The shape of the marker differentiates stimulus pairs with different relative stimulus distance and therefore different difficulty, from 8 mm (triangle) to 40 mm (circle). Colors indicate the percentage of the S2-ward bias. (a) Mean performance variations for trials with low values of the previous S2, i.e. 8 or 16 mm. Similarly to what is shown in Fig. 2a the prevalence of red markers indicates a general positive S2-ward bias (similarly to what is shown in Fig. 2a left side, but here extended to all stimulus pairs). (b) Mean performance variations for trials with high values of previous S2, i.e. 40 or 48 mm. In line with Fig. 2a right side, the prevalence in this case of blue markers reveals a negative S2-ward bias.

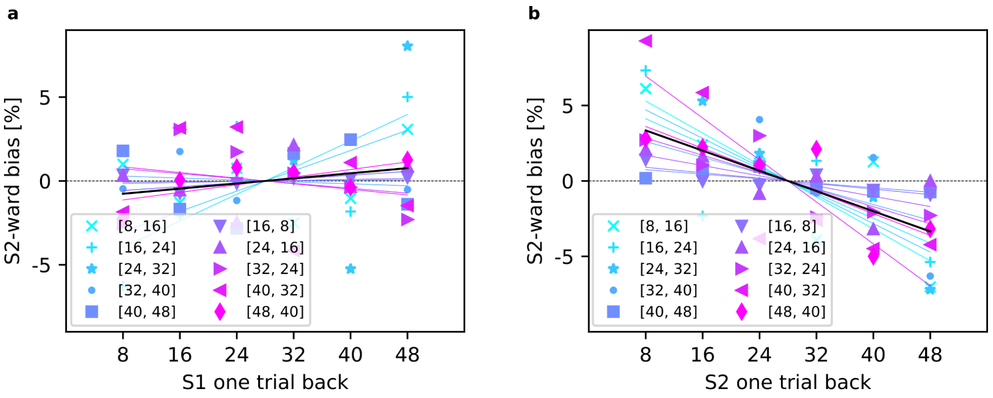


Figure S2. Effect of the previous S1 and S2 stimuli on the current response. (a) Effect of the previous S1 on the current response. (b) Effect of the previous S2 on the response in the current trial (this last panel is the same as Fig. 2a, reported here just for comparison). Comparing (a) with (b) it appears that only the previous S2 and not the previous S1 affects the performance (a: linear fit slope=0.31, p=0.14; b: linear fit slope=-1.34, p<0.0001). This comparison is important to exclude a repeat/shift response bias.

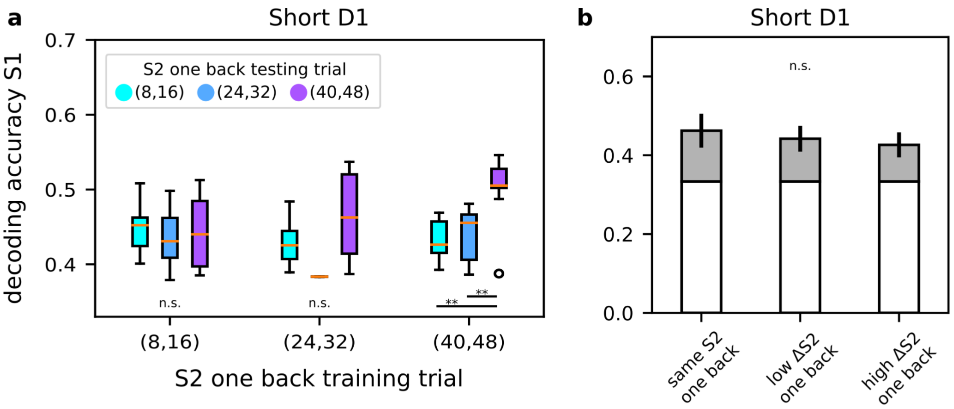


Figure S3. Decoding accuracy of stimulus S1 conditioned on S2 of the previous trial, boxplots show how the accuracy changes in the first 400 ms of delay 1 (short D1) panel (a) when training and testing trials share (or not) their S2 in one-trial back. n.s.=not significant, *=p<0.05, **=p<0.01, ANOVA test with Tukey's multiple comparison test. (b) Same results as in (a), but grouped in: decoding of S1 within condition (same S2 previous trial) and across conditions (low ΔS2: small gap between S2 one-trial back in train and test sets, e.g. (8,16) vs. (24,32), high ΔS2: large gap between S2 one-trial back in train and test sets, e.g. (8,16) vs. (40,48)). n.s.=not significant, ANOVA test with Tukey's multiple comparison test.

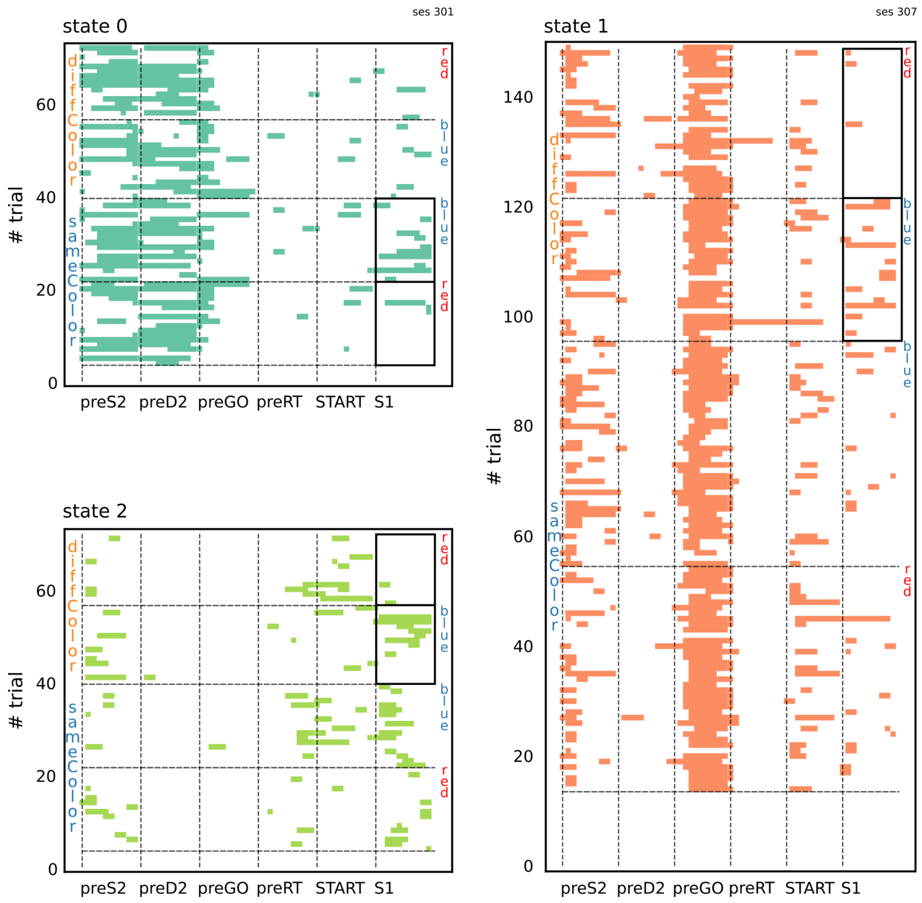


Figure S4. Examples of sessions with significant HMM coding states for the first stimulus color feature. This figure extends Fig. 4a by presenting the HMM state sequences for each coding state at the single-trial level, spanning from the appearance of S2 in the previous trial (preS2) to the end of S1 presentation in the current trial. Trials are grouped by the diffColor or sameColor condition, and within each condition, by the stimulus visual feature (blue or red). The presence of coding states is highlighted with black bold rectangles, which enclose subsets of trials where the mean state occupancy significantly differs between different stimulus visual features (Mann-Whitney U test with Benjamini/Hochberg false discovery rate correction, $p<0.05$).
